# Hi-C yields chromosome-length scaffolds for a legume genome, *Trifolium subterraneum*

**DOI:** 10.1101/473553

**Authors:** Olga Dudchenko, Melanie Pham, Christopher Lui, Sanjit S. Batra, Marie Hoeger, Sarah K. Nyquist, Neva C. Durand, Muhammad S. Shamim, Ido Machol, William Erskine, Erez Lieberman Aiden, Parwinder Kaur

## Abstract

We present a chromosome-length assembly of the genome of subterranean clover, *Trifolium subterraneum*, a key Australian pasture legume. Specifically, *in situ* Hi-C data (48X) was used to correct misjoins and anchor, order, and orient scaffolds in a previously published genome assembly (TSUd_r1.1; scaffold N50: 287kb). This resulted in an improved genome assembly (TrSub3; scaffold N50: 56Mb) containing eight chromosome-length scaffolds that span 95% of the sequenced bases in the input assembly.

## Introduction

Sustainable agricultural production entails growing food without damaging the underlying soil (Tilman et al. 2011; Long, Marshall-Colon, and Zhu 2015). Legumes are of great interest for such efforts: because they produce their own nitrogen via symbiotic nitrogen fixation, legumes can actually improve the soil (Chikowo et al. 2004; Saha, and Mandal 2009). Among legumes, pasture legumes tend to be more resilient to stress and more capable of thriving in marginal land (Beuselinck et al. 1994; Nichols et al. 2013; Hirakawa, Kaur et al. 2016).

Two years ago, we published a draft genome of subterranean clover, *Trifolium subterraneum* (Hirakawa, Kaur et al. 2016). The draft, TSUd_r1.1, was created using a combination of Illumina and Roche 454 GS FLX+ sequencing. Single-end, 520-660bp paired-end and 2kb, 5kb, 8kb, 10kb, 15kb and 20kb mate-pair libraries were constructed, generating 8297 scaffolds larger than 1kb, with contig N50 of 22kb and scaffold N50 of 287kb and spanning 403Mb of sequence.

Recently, we and others have shown that it is possible to significantly improve draft genomes by using data derived from *in situ* Hi-C (Lieberman-Aiden, van Berkum et al. 2009; Rao, Huntley et al. 2014; Burton et al. 2013; Dudchenko et al. 2017). Because Hi-C can estimate the relative proximity of loci in the nucleus, Hi-C contact maps can be used to correct misjoins, anchor, order, and orient contigs and scaffolds. This process greatly improves contig accuracy and typically yields chromosome-length scaffolds.

To broaden the range of genetic resources available for legumes, we used Hi-C to improve the TSUd_r1.1 draft assembly, producing a genome assembly for *Trifolium subterraneum* with chromosome-length scaffolds.

## Results

### An assembly of Trifolium subterraneum with chromosome-length scaffolds

We began by generating *in situ* Hi-C data (Lieberman-Aiden, van Berkum et al. 2009; Rao, Huntley et al. 2014) from *Trifolium subterraneum* leaves.

We then improved the TSUd_r1.1 using the approach described in (Dudchenko et al. 2017, 2018). First, we set aside scaffolds shorter than 10kb, leaving 3161 scaffolds. Next, we ran the 3D De Novo Assembly (3D-DNA) pipeline using our Hi-C data in order to anchor, order, orient, and correct misjoins in the TSUd_r1.1 scaffolds (see Fig. 1) (Dudchenko et al. 2017). Finally, we performed a manual refinement step using Juicebox Assembly Tools (Dudchenko et al. 2018). The resulting assembly, dubbed TrSub3, comprises 8 chromosome-length scaffolds, whose lengths range from 49.5Mb to 65.2Mb. These chromosome-length scaffolds span 99.6% of the sequenced bases in the 3161 input scaffolds. The remaining 0.4% of the sequence is included in 347 small scaffolds (scaffold N50: 25kb). See Tables 1, S1-S5.

**Table 1:** Assembly statistics for TrSub3 genome assembly. Note that scaffolds smaller than 1000 base pairs are excluded from the analysis.

| **TrSub3** |  |
| --- | --- |
| **Draft scaffolds** |  |
| **Base Pairs** | 403,400,476 |
| **Number of contigs** | 46,474 |
| **Contig N50** | 22,377 |
| **Number of scaffolds** | 8,297 |
| **Scaffold N50** | 286,571 |
| **Chromosome-length scaffolds** |  |
| **Base Pairs** | 392,723,996 |
| **Number of contigs** | 39,734 |
| **Contig N50** | 23,051 |
| **Number of scaffolds** | 8 |
| **Scaffold N50\*** | 56,309,329 |
| **Small scaffolds** |  |
| **Base Pairs** | 1,949,837 |
| **Number of contigs** | 834 |
| **Contig N50** | 9,493 |
| **Number of scaffolds** | 347 |
| **Scaffold N50** | 24,672 |
| **Tiny scaffolds** |  |
| **Base Pairs** | 8,726,933 |
| **Number of contigs** | 5,867 |
| **Contig N50** | 1,529 |
| **Number of scaffolds** | 4,930 |
| **Scaffold N50** | 2,883 |

**Figure 1:**
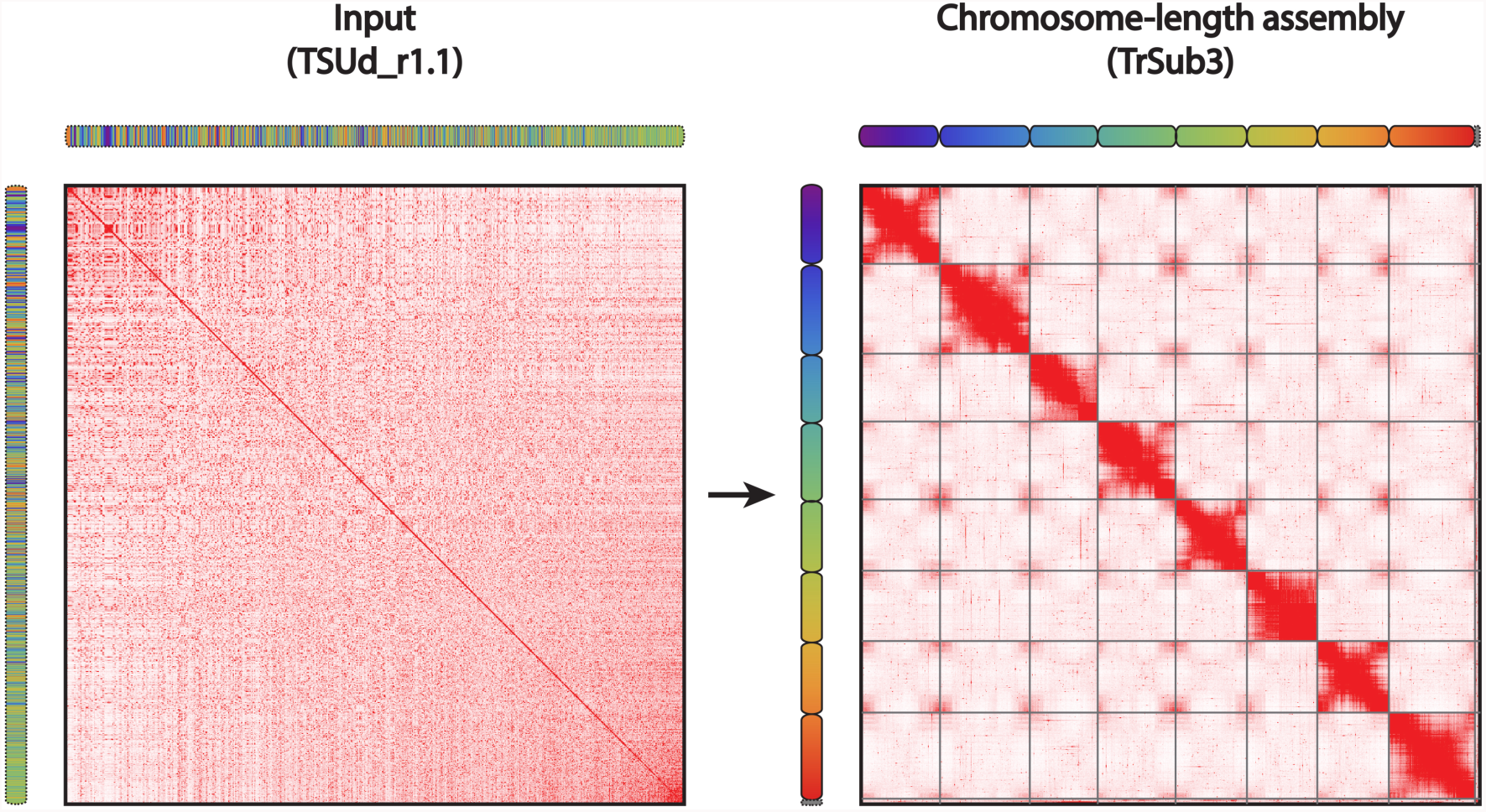
Hi-C map of the draft and chromosome-length assemblies of *Trifolium subterraneum* genome. Contact matrices were generated by aligning the same Hi-C data set to TSUd_r1.1 draft genome (left) and the TrSub3 genome assembly generated using Hi-C (right). Pixel intensity in the matrix indicates how often a pair of loci collocate in the nucleus. The correspondence between loci in the draft and final assemblies is illustrated using chromograms. The chromosome-scale assembly scaffolds in TrSub3 are assigned a linear color gradient, and the same colors are then used for the corresponding loci in the TSUd_r1.1 (left). The draft scaffolds are ordered by sequence name. Grid lines highlight the boundaries of eight chromosome-length scaffolds in TrSub3 (right). Scaffolds smaller than 10kb in TSUd_r1.1 are not included in this illustration.

### High degree of synteny and collinearity between the subterranean clover and the barrel clover

The availability of a genome assembly with chromosome-length scaffolds for the closely related model legume barrel clover (*Medicago truncatula)*, MedtrA17_4.0, enabled us to study the evolution of clover genomes. We performed a whole-genome alignment between these two species using LastZ (Robert S. Harris 2007)^1^. We found extensive synteny, in the sense that loci on the same chromosome in *T. subterraneum* tend to lie on the same chromosome in *M. truncatula.* We also observed chromosome-scale collinearity of the barrelclover and subterranean clover chromosomes, in the sense that loci tend to appear in the same linear order.

We compared these results with those obtained in a prior study, where we used optical and linkage maps to improve TSUd_r1.1, and to thereby generate large scaffolds for *T. subterraneum* (Hirakawa, Kaur et al. 2016; Kaur et al. 2017). (Note that these optical and linkage maps were not employed in the generation of TrSub3.) Like the present study, the previous study, whose assembly was dubbed Tsub_Refv2.0, reported that loci on the same chromosome in *T. subterraneum* tend to lie on the same chromosome in MedtrA17_4.0. However, several chromosomes in Tsub_Refv2.0 did not exhibit large blocks of collinearity with *M. truncatula* (Kaur et al. 2017). Instead, the order of the loci on these chromosomes was extensively permuted between the two species. This comparison suggests that the chromosome-length scaffolds in the TrSub3 genome are more consistent with those of the MedtrA17_4.0 genome assembly.

## Discussion

Several limitations of the TrSub3 assembly reported here ought to be borne in mind. First, errors in the input assembly may remain in the final genome. The most common scenario is likely to be small insertions in the draft scaffolds that are not identified by our misjoin detection algorithm (see Dudchenko et al. 2017). Second, about 5% of the sequence reported in TSUd_r1.1 (3% of sequence in scaffolds larger than 1kb) is not anchored in TrSub3. Instead, these sequences are partitioned among multiple small scaffolds that have relatively low Hi-C coverage and are thus more difficult to analyze. Finally, the current approach is not perfect for local ordering of very small adjacent contigs (see Dudchenko et al. 2017). Despite these limitations, comparative analysis with other legume species (see Fig. 2) suggests that TrSub3 is a considerable improvement over prior efforts to advance subterranean clover assembly using genetic and optical mapping (Hirakawa, Kaur et al. 2016; Kaur et al. 2017).

**Figure 2:**
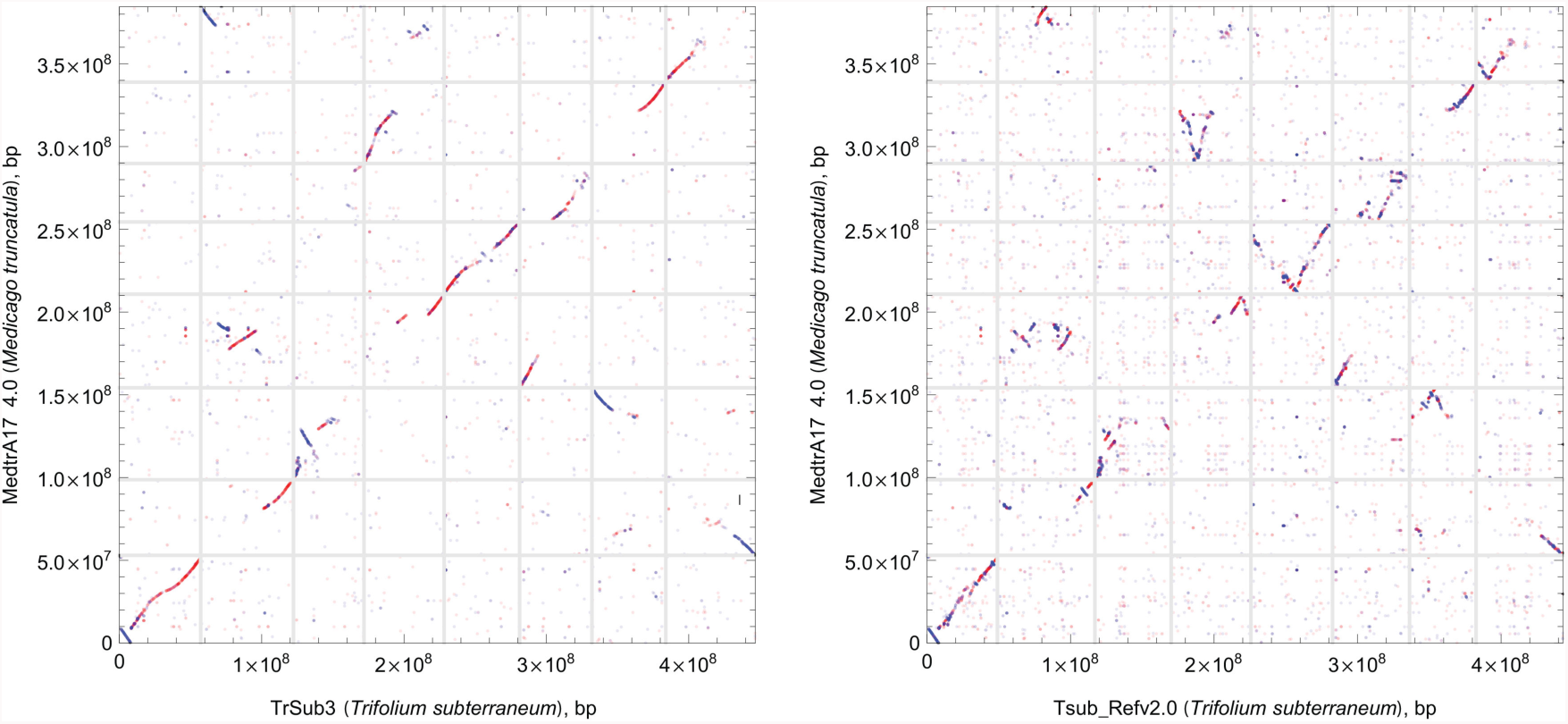
Assembly using Hi-C improves comparative analysis. The analysis of synteny between *Medicago truncatula* and *Trifolium subterraneum* suggests that the new assembly (left, TrSub3) better reflects the large-scale structure of the genome than recently published scaffolds assembled using optical and linkage mapping (right, Tsub_Refv2.0). For this analysis, the *T. subterraneum* and *M. truncatula* fastas were aligned using the LastZ alignment algorithm (Robert S. Harris 2007). Here, we show alignment blocks with scores larger than 50,000 (Robert S. Harris 2007), with direct synteny blocks colored red, and inverted blocks colored blue.

It is worth noting that the Hi-C assembly approach also yields insight into the three-dimensional structure of the subterranean clover genome (see Fig. 1). The latter has the potential to broaden our understanding of gene regulation including genes associated with abiotic stress tolerance and legume nodulation as well as increase our ability to manipulate plant genomes. We hope that the new assembly and data generated will be of service in resolving food insecurity as well as sustainable soil improvement.

## Materials and Methods

In situ Hi-C was performed as described previously (Rao, Huntley et al., 2014) using fresh leaves from subterranean clover (*T. subterraneum*) cv. Daliak. Prior to harvesting, mature dry seeds were grown for 2-3 weeks in sterilized potting mix and dark treated for 2-3 days. The resulting library was sequenced to yield approximately 48X coverage of the *T. subterraneum* genome. The library was processed against TSUd_r1.1 using the Juicer pipeline (Durand, Shamim, et al. 2016) and assembled following the methods described in (Dudchenko et al. 2017, 2018). The resulting contact maps were visualized using 3D-DNA and Juicebox visualization system (Durand, Robinson, et al. 2016; Dudchenko et al. 2017, 2018).

## Acknowledgements

Meat & Livestock Australia and Centre for Plant Genetics and Breeding (PGB) provided financial support to the study. We acknowledge the supercomputing resources provided by the Pawsey Supercomputing Centre with funding from the Australian Government and the Government of Western Australia. This work was supported by a Center for Theoretical Biological Physics postdoctoral fellowship to O.D., an NIH New Innovator Award (1DP2OD008540-01), an NSF Physics Frontiers Center Award (PHY-1427654, Center for Theoretical Biological Physics), the Welch Foundation (Q-1866), an NVIDIA Research Center Award, an IBM University Challenge Award, a Google Research Award, a Cancer Prevention Research Institute of Texas Scholar Award (R1304), a McNair Medical Institute Scholar Award, an NIH 4D Nucleome Grant U01HL130010, an NIH Encyclopedia of DNA Elements (ENCODE) Mapping Center Award UM1HG009375, the President’s Early Career Award in Science and Engineering to E.L.A.

## Author Contributions

P.K., O.D. and E.L.A conceived the project. M.P. and C.L. adapted the Hi-C experiment to plant tissue and performed the Hi-C experiments. O.D., M.P., C.L., S.S.B., M.H., S.K.N., N.C.D., M.S.S., I.M. and E.L.A. analyzed the data. P.K., O.D., W.E. and E.L.A. wrote the manuscript with contributions from all authors.

## Accession code

Chromosome-length genome sequence assembly (TrSub3) and annotation data has been made available at Trifolium GBrowse Webpage http://trifoligate.info/.

## Restrictions on use

The TrifoliGATE Expert Working Group makes the TrSub3 genome assembly available to the research community under the agreement that as producers of the chromosome-scale assembly, we reserve the right to be the first to publish a genome-wide analysis incorporating this assembly. Researchers are encouraged to contact us if there are queries about referencing or publishing analyses based on the data described in the manuscript. Researchers are also invited to consider collaborations with the TrifoliGATE Expert Working Group for larger studies or if the limitations here restrict further work.

## Supplementary Data

**Table S1:** Statistics describing various scaffold populations. See (Dudchenko et al. 2017). Scaffolds smaller than 1000 base pairs are excluded from the analysis.

| Scaffold Type | Statistic | TrSub3 |
| --- | --- | --- |
| Draft | Sequenced Base Pairs | 403,400,476 |
|  | # of Scaffolds | 8,297 |
|  | Scaffold N50, bp | 286,571 |
|  | Length of Longest Scaffold, bp | 2,878,652 |
| Unattempted | Sequenced Base Pairs | 8,994,298 |
|  | % of Total Sequenced Base Pairs | 2.2% |
|  | # of Scaffolds | 5,136 |
|  | Scaffold N50, bp | 2,872 |
|  | Length of Longest Scaffold, bp | 9,989 |
| Input | Sequenced Base Pairs | 394,406,178 |
|  | % of Total Sequenced Base Pairs | 97.8% |
|  | # of Scaffolds | 3,161 |
|  | Scaffold N50, bp | 294,854 |
|  | Length of Longest Scaffold, bp | 2,878,652 |
| Resolved | Sequenced Base Pairs | 392,723,996 |
|  | % of Attempted | 99.6% |
|  | % of Total Sequenced Base Pairs | 97.4% |
|  | # of Scaffolds | 2,988 |
|  | Scaffold N50, bp | 299,815 |
|  | Length of Longest Scaffold, bp | 2,878,652 |
| Unresolved & Inconsistent | Sequenced Base Pairs | 1,949,837 |
|  | % of Attempted | 0.49% |
|  | % of Total Sequenced Base Pairs | 0.48% |
|  | # of Scaffolds | 383 |
|  | Scaffold N50, bp | 20,037 |
|  | Length of Longest Scaffold, bp | 319,620 |

**Table S2:**
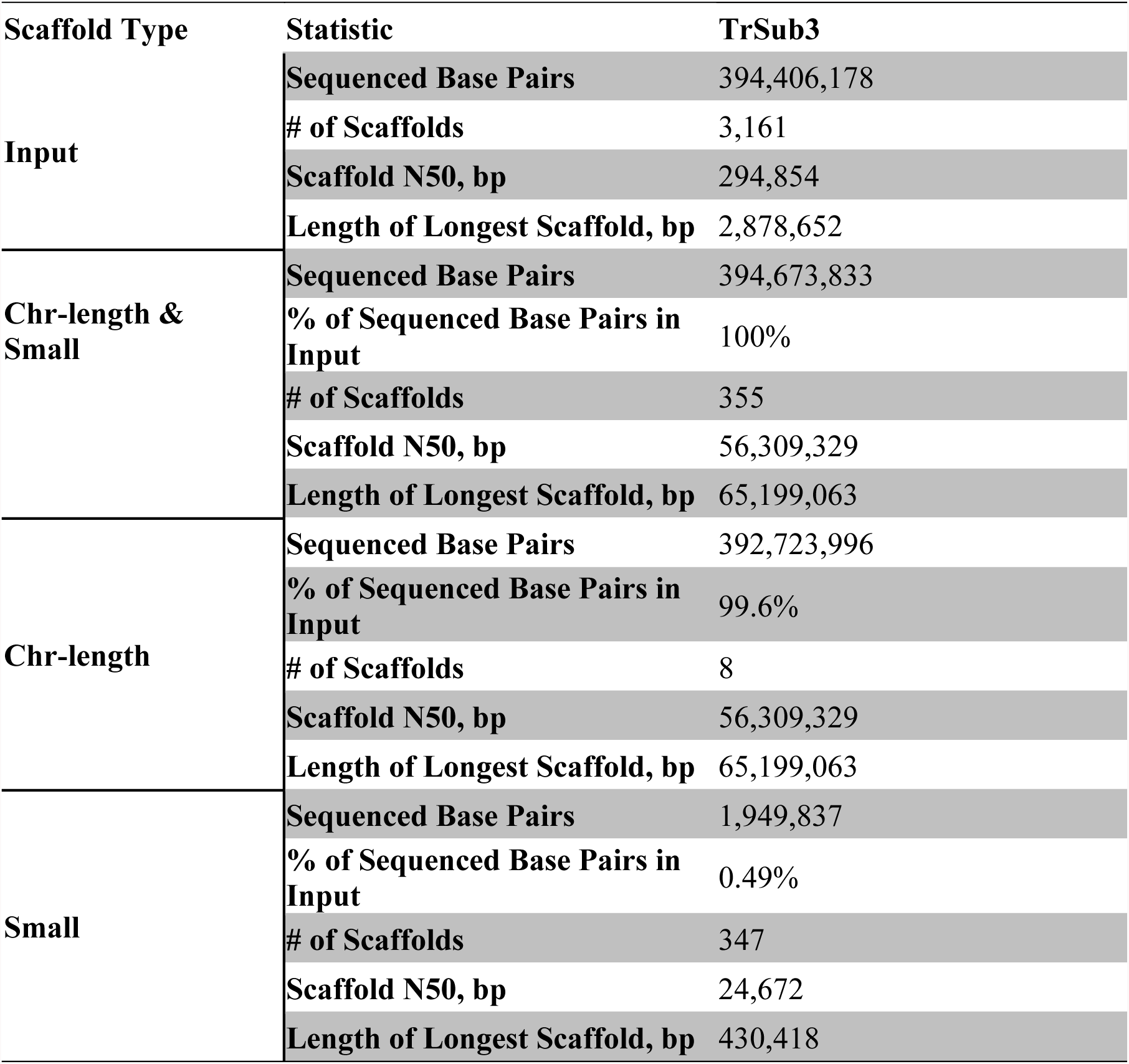
Statistics describing the results of assembly using Hi-C, including only draft scaffolds that we attempted to scaffold further as input. The values describe the chromosome-length scaffolds, as well as other, smaller scaffolds generated during the Hi-C assembly process.

**Table S3:** Cumulative assembly statistics for the assemblies. The values describe the combined set of chromosome-length scaffolds, as well as small scaffolds; however, they exclude the ‘tiny’ scaffolds (<10kb) from the draft assembly, which we did not attempt to assemble.

| **Statistics** | **TrSub3** |
| --- | --- |
| **Base Pairs** | 394,673,833 |
| **Number of contigs** | 40,568 |
| **Contig N50** | 22,999 |
| **Number of scaffolds** | 355 |
| **Scaffold N50** | 56,309,329 |
| **In chromosome-length scaffolds** | 99.5% |

**Table S4:** Cumulative assembly statistics for the assemblies. The values describe the combined set of chromosome-length scaffolds, as well as small and tiny scaffolds; ‘subtiny’ scaffolds (<1kb) from the draft assembly are excluded from this analysis.

| **Statistics** | **TrSub3** |
| --- | --- |
| **Base Pairs** | 403,400,766 |
| **Number of contigs** | 46,435 |
| **Contig N50** | 22,377 |
| **Number of scaffolds** | 5,285 |
| **Scaffold N50** | 56,309,329 |
| **In chromosome-length scaffolds** | 97.4% |

**Table S5:** Chromosome-length scaffolds of TrSub3 genome assembly.

| **Chr-length scaffold** | **Total length, bp** | **Sequenced bases, bp** |
| --- | --- | --- |
| **1** | 57,151,827 | 51,189,519 |
| **2** | 65,199,063 | 57,810,030 |
| **3** | 49,527,580 | 42,686,100 |
| **4** | 56,309,329 | 50,380,603 |
| **5** | 53,036,926 | 46,854,310 |
| **6** | 50,764,665 | 43,812,303 |
| **7** | 51,904,178 | 46,132,021 |
| **8** | 62,889,439 | 53,840,477 |

## Footnotes

1 The alignment was run as follows: lastz target query --notransition --step=20 –-nogapped. No chaining was performed.

